# Aligning Needs: Integrating Citizen Science Efforts into Schools Through Service Requirements

**DOI:** 10.1101/304766

**Authors:** Ginger Tsueng, Arun Kumar, Steven M. Nanis, Andrew I Su

## Abstract

Citizen science is the participation in scientific research by members of the public, and it is an increasingly valuable tool for both scientists and educators. For researchers, citizen science is a means of more quickly investigating questions which would otherwise be time-consuming and costly to study. For educators, citizen science offers a means to engage students in actual research and improve learning outcomes. Since most citizen science projects are usually designed with research goals in mind, many lack the necessary educator materials for successful integration in a formal science education (FSE) setting. In an ideal world, researchers and educators would build the necessary materials together; however, many researchers lack the time, resources, and networks to create these materials early on in the life of a citizen science project. For resource-poor projects, we propose an intermediate entry point for recruiting from the educational setting: community service or service learning requirements (CSSLRs). Many schools require students to participate in community service or service learning activities in order to graduate. When implemented well, CSSLRs provide students with growth and development opportunities outside the classroom while contributing to the community and other worthwhile causes. However, CSSLRs take time, resources, and effort to implement well. Just as citizen science projects need to establish relationships to transition well into formal science education, schools need to cultivate relationships with community service organizations. Students and educators at schools with CSSLRs where implementation is still a work in progress may be left with a burdensome requirement and inadequate support. With the help of a volunteer fulfilling a CSSLR, we investigated the number of students impacted by CSSLRs set at different levels of government and explored the qualifications needed for citizen science projects to fulfill CSSLRs by examining the explicitly-stated justifications for having CSSLRs, surveying how CSSLRs are verified, and using these qualifications to demonstrate how an online citizen science project, Mark2Cure, could use this information to meet the needs of students fulfilling CSSLRs.

## 1. Introduction

Community service is a voluntary non-curriculum-based activity that may be recognized (and in some cases, required) by schools and generally does not have explicit learning objectives. In contrast, service learning is a curriculum-based community service activity that has explicit learning objectives (Spring, Grimm Jr, and Dietz, 2008). For example, participating in a beach clean-up would be considered a community service activity; however, if the activity was part of a class curricula on the pollution and its effects on the food web, it would be considered a service learning activity. When implemented well, service learning has been associated with improvements in cognitive skills/academic performance, civic engagement, social skills, and improved attitudes towards the self and towards school and learning (Celio, Durlak, and Dymnicki, 2011)(McLellan and Youniss, 2003)(Shumer, 1994). Because of these benefits, twenty nine states have policies in place to encourage community service or service learning at the high school level, but only one state (Maryland) has a statewide community service or service learning requirement (CSSLR) (Ecs.force.com, 2014). Even without a state-set CSSLR, cities like Chicago and Washington D.C. have city-wide CSSLRs, and in the absence of city-set CSSLRs, CSSLRs may be imposed by school districts or individual schools. In Canada, the province of Ontario has a province-wide CSSLR.

Once mandated, the impact of community service on student development becomes highly dependent on the way the CSSLRs are explained and implemented. For example, participation in community service activities in high school has been associated with increased likelihood for participating in community service activities after graduation even when the community service was mandated (Metz and Youniss, 2003). However, this increased propensity may be restricted to students who felt they maintained some autonomy in spite of the requirements (Stukas, Snyder and Clary, 1999), and the observed improvement in volunteer rates disappear eight years after high school graduation (Planty and Regnier, 2003). Student perceptions on CSSLRs depend on how schools convey the requirements to students. When perceived as a burdensome requirement for graduation, students reported deriving less value out of their CSSLR-fulfilling activities, especially if no clarity is provided on why a CSSLR-fulfilling activity meets the requirement (Jones, Segar and Gasiorski, 2008). Successful implementation of CSSLRs requires careful planning, deliberate articulation of rationale and expectations, resources, community relationships, and iterative review and improvement of the program (Raskoff and Sundeen, 1999). Without sufficient infrastructure and resources in place to properly implement CSSLRs, CSSLRs may encumber students and educators with record keeping, transportation, and scheduling burdens that can ultimately negate the positive benefits describe in the literature (Loup, 2000).

Participation in community service and service learning activities has been found to be particularly beneficial for students at risk of academic failure (Schmidt, Shumow, and Kackar, 2012) yet students from lower socioeconomic status, who stand to gain the most from community service / service learning activities are much less likely to engage in them (Planty and Regnier, 2003). By having CSSLRs in place, schools may address the motivational barrier by forcing participation, and if implemented well, schools may also address a key logistical barrier by helping students identify opportunities for participation (Nolin et al., 1997) (Hart, Atkins, and Ford, 1998). However, many of the same barriers to participation in sports activities also exist for community service activities and will further burden students if schools have CSSLRs in place without additional plans/assistance in place. These barriers include: time (parent’s schedule, student’s schedule, transportation time), cost (transportation costs, equipment costs), and location (access and transportation)(Raskoff and Sundeen, 1999)(Somerset and Hoare, 2018). In schools where CSSLRs were set without sufficient administrative, educator, and student support/resources, students–especially those of lower socioeconomic status–may find the CSSLR itself to be a barrier to graduation, and educators may find themselves creating tasks to help students fulfill CSSLRs (Jones, Segar and Gasiorski, 2008). For these students and educators, we propose a potential means of fulfilling CSSLRs– citizen science projects.

Citizen science is a growing practice which involves inviting anyone with an interest in science to contribute to scientific endeavors, and has become an increasingly valuable tool for scientists and educators alike (Bonney et al., 2009). For researchers, citizen science offers a way to collect granular data from broad geographic areas over time, sort and analyze large datasets quickly, and ultimately expand scientific knowledge (Raddick et al., 2009). For educators, citizen science presents a unique opportunity to involve students in ongoing scientific endeavors (Haines, n.d.) and provides a forum for engaging in scientific inquiry (Trumbull et al., 2000). Although many projects were not originally designed with an educational goal in mind and educational support in citizen science projects remains highly variable, limited informal learning has been observed in association with citizen science (Brossard et al., 2005)(Masters et al. 2016). Furthermore, the high potential value of applying citizen science towards informal science education warranted the development of a framework which aligns the needs of stakeholders from citizen science and informal science education (ISE) (Shirk et al, 2012).

Similar to community service activities, many citizen science projects have a diversity problem when it comes to the volunteers they are able to recruit (Land-Zandstra et al., 2015). Environmental volunteers are less likely to come from lower socioeconomic status or be of ethnic minorities backgrounds. In the interest of reaching and engaging a wider and more inclusive audience in science (democratizing science) and fulfilling their own recruitment needs, citizen science researchers are looking for ways to successfully incorporate citizen science into formal science education (Shah and Martinez, 2016) (He and Wiggins, 2017). The administrative burden for incorporating citizen science into formal science education can be high considering entry barriers such as: limitations in project resources (He and Wiggins, 2017); dependence on technological availability (Mueller, Tippins, and Bryan, 2012) (Silva et al., 2016); and limits in educator access to scientific materials (Gray, Nicosia, and Jordan, 2012), expertise (Bonney et al., 2015) and time (Silva et al., 2016). In spite of these barriers, citizen science projects have been successfully integrated into the classroom setting (Butler and MacGregor, 2003)(Gray, Nicosia, and Jordan, 2012). As with service learning implementation, successful integration of citizen science projects in the classroom requires meaningful interactions between educators and collaborators (in this case scientists) to resolve differences in stakeholder priorities (Zoellick, Nelson, and Schauffler, 2012) and the generation of educational resources. Scientists need to work with educators to create educational resources, yet many educators are under severe time constraints which makes them difficult to recruit without already having educational resources in place. To bypass this chicken-and-egg problem of educator and educational material, citizen science projects can utilize CSSLRs as a third entry point into the educational environment.

Citizen science projects, especially virtual citizen science projects, can potentially be used to meet the needs of students facing CSSLRs who do not have sufficient support for overcoming time, cost, and location barriers. In particular, students with scheduling difficulties due to part-time jobs or familial obligations would be able to fulfill their CSSLRs with accordance to their own schedule. With virtual citizen science projects, students would be able to fulfill CSSLR obligations from home or school, effectively bypassing transportations barriers associated with fulfilling CSSLRs in person. To investigate the use of citizen science projects for fulfilling CSSLRs, we first investigate CSSLRs and enrollment numbers in San Diego County Public High Schools to estimate the number of students impacted by CSSLRs and compare them with estimates from regions with known citywide, statewide, or provincewide CSSLRs. Surveys on CSSLRs in the literature tend to directly target students or educators and provide deep and meaningful insight on the CSSLRs their relationship to CSSL activities (Nolin et al., 1997)(Raskoff and Sundeen, 1999), but often do not provide a general estimate of the number students impacted by CSSLRs in the surveyed areas.

With the help of a volunteer fulfilling service requirements we inspect how CSSLRs are explicitly conveyed in San Diego County. Since student experience is affected by their perception of the requirements (Jones, Segar and Gasiorski, 2008), it is important to examine the rationales expressed by schools that encourage and/or require CSSL activities. We also identify the kinds of activities that are qualified to fulfill CSSLRs and how CSSLRs fulfillments are verified as understanding these factors will allow citizen science projects to align their opportunities with the needs of the students seeking to fulfill CSSLRs. Assuming that most community service activities aim to benefit people, we next review the citizen science projects listed in SciStarter as health/medicine citizen science projects (because health/medicine ultimately impact people) to broadly understand the current alignment between factors of CSSLRs and citizen science. Lastly, we use an example, Mark2Cure, to demonstrate how these factors may be used to better align an online citizen science project towards fulfilling CSSLRs.

This paper is organized as follows: After this introductory section, Section 2 describes the methods used to investigate the landscape of CSSLR (i,e.-the potential need generated by CSSLRs, the qualifications of a CSSL activity, the verification methods for CSSLRs), and the suitability of citizen science projects for fulfilling CSSLRs (i.e.-the identification of citizen science projects that can potentially fulfill a CSSLR, and how CSSLRs may be fulfilled with a virtual citizen science project, Mark2Cure). Section 3 describes the results and findings and ends with recommendations based on the those findings. Section 4 describes the role of citizen science in the formal and informal science education setting, the role of volunteerism in education, and discusses the barriers generated by CSSLRs and technology access for students from lower socioeconomic status. Section 5 concludes the paper.

## 2. Methods

### 2.1 Overview of the high school CSSLR landscape at different levels of government

To roughly estimate of the market size of students in with CSSLRs, we investigate the number of students at different geographic levels of CSSLRs. There are a limited number of examples of high school CSSLRs set outside the individual school or school district level. At the city-wide level, Chicago and Washington DC are examples of cities with CSSLRs. As seen in Table 1, we pulled data on Chicago’s high school CSSLRs from the Chicago Public Schools Policy Manual (Chicago Board of Education, 2017). At the state level, Maryland is the only state to have statewide high school CSSLRs in the US, and information on Maryland CSSLRs. There are no other statewide high school CSSLRs in the US, but Canada provides an example of provincewide high school CSSLRs. High school enrollment data were obtained as listed in Table 1. Data on community service participation was available for Maryland, but had to be estimated for Chicago and Ontario by averaging the requirement over the four years expected for high school students and multiplying by the number of high school students.

**Table 1.** Summary of data sources used to derive the overview

| Geographic level | Example | Data type | Reference |
| --- | --- | --- | --- |
| Citywide | Chicago | CSSLR | Chicago Board of Education, 2017 |
| Citywide | Chicago | Enrollment | Cps.edu, 2018 |
| Statewide | Maryland | CSSLR | Code of Maryland Regulations, 2016 |
| Statewide | Maryland | Enrollment | Mdp.state.md.us, 2016 |
| Statewide | Maryland | Participation | Marylandpublicschools.org, 2017 |
| Provincewide | Ontario | CSSLR | Ontario Ministry of Education, 2016 |
| Provincewide | Ontario | Enrollment | Ontario Ministry of Education, 2018 |

### 2.2 Surveying the high school CSSLR landscape in San Diego

A list of public San Diego high school districts, high schools, and the 2016-2017 student enrollments at each school were obtained from the California Department of Education (Ed-data.org, 2018). CSSLR requirements for each school were investigated via school or district-specific website searches and inquiries by phone or email. For schools with no CSSLRs, we investigated Junior Reserve Officer Training Corps (JROTC) programs and/or Key clubs (Kiwanis.org-associated volunteering club), both of which tend to have community service requirements. Key club membership estimates were based on the Key Club International Paid Clubs Report for the 2016-2017 school year available from the Kiwanis International website (Kiwanis.org, 2017). These numbers account for only due-paying members and may underestimate the actual membership since payment of dues is not needed for participation and volunteer activities in some schools.

### 2.3 Estimating students impacted by or engaging in CSSLRs/CSSL activities in San Diego

Rough estimates of students engaged in CSSLRs were obtained as follows:

1. If a school has a CSSLR, the number of students impacted by the CSSLR was assumed to be 100% and therefore the number of students enrolled in that school was used.
2. If a school does NOT have CSSLRs, any estimation on student participation in volunteer activities provided by the schools were used.
3. If a school does NOT have CSSLRs or estimates on student participation in volunteer activities, the school was checked for the presence of Key Club or a JROTC program. The memberships in both programs were estimated and the number of volunteers was assumed to be the higher membership of the club/program.
4. If a school does not have CSSLRs, estimates on student participation, key club or JROTC, it was treated as not estimateable.

### 2.4 Qualitative inspection of the rationale for CSSLRs and community service recommendations or mentions, verification methods, and qualifying activities

The websites and student/parent handbooks (publicly available materials) of all schools in this study were expected to be highly region-specific, subject to change, but still informative. For this reason, they were inspected for any mention of community service by a single reviewer. If community service was mentioned on the school’s publicly available materials, the content was inspected for language that encouraged or recommended engaging in community service. If the school mentioned, endorsed or recommended engaging in community service:

1. The expressed reasons/rationale given for the mention/endorsement were qualitatively binned into thematic categories with new categories added whenever a rationale did not fit in with an existing one. These categories were then further binned into the five service program educational goals established by Furco (1994). An example of the qualitative binning process can be found in Supplementary Figure 1. The intermediate list of categories along with the corresponding Furco educational goals is as follows:

a. Admissions: Because college admissions (academic achievement)
b. Connection: Foster civic responsibility and community/engagement, relationships (social development, political development)
c. Skills: Learn/build skills (academic achievement, vocational development)
d. Humanitarian: Improve community/school/help neighbors (social development)
e. Character: Build esteem, good feels, character (personal and moral development)
f. Experience: Real life experience (vocational development)
g. Merit: Qualify for award, association, guaranteed admission, IB qualification etc. (academic achievement)
h. Fun: For fun/interest/passion (personal and moral development)
i. Requirement: Particular/specific class requirement (academic achievement)
j. Intervention: Intervention / penalty (political development)
2. The school’s website was also searched for the availability of a sample tracker, log, form, or other guidelines for verifying a student’s participation in community service activities.

a. If a tracker, log, form or other guideline could be found, the fields on that tracker, log, or form were documented.
3. The school’s website was checked for guidelines on activities which qualified vs didn’t qualify as community service activities, and the sample activities were recorded and binned.

### 2.5 Reviewing health and medical citizen science projects for factors which affect student fulfillment of CSSLRs

Scistarter is a portal that facilitates interaction between different stakeholders of citizen science projects (Cavalier et al., n.d.). Project leaders can add their projects to SciStarter and thematically categorize their efforts. Assuming that at minimum, community service requirements exist to help other people, we filtered for Citizen Science projects in health and medicine since those would logically have a very direct impact on people. There were 101 projects that were identified as health and medical citizen science projects and we inspected these projects for the organization type, non-profit status, frequency/availability of the opportunity, cost to participate, participant role, and whether or not contributions can be done from home. The organization behind the project, the cost to participate, frequency/availability of the opportunity, and whether or not contributions can be done from home were self-reported by the projects to SciStarter. The organizations behind the project were classified for their for-profit/non-profit status based on US tax definitions for non-profit organizations. Non-profit organizations were further categorized based on types of 501(c) organizations with government agency as an additional type.

### 2.6 Aligning Mark2Cure metrics with CSSLRs

Mark2Cure is a virtual citizen science project that enables volunteers to extract information buried in biomedical abstracts (Tsueng et al., 2016). With the help of volunteers, Mark2Cure aims to organize knowledge to facilitate biomedical discovery. It is currently focused on biomedicals in the NGLY1-deficiency rare disease space. Volunteers complete and online tutorial which teaches them how to use the system to perform Named Entity Recognition (NER)–the first step in information extraction. We collected annotations from Mark2Cure’s NER missions completed between 2015.05.01 and 2017.11.22. Annotations that were marked by at least 6 users were used as the reference set. Abstracts in Mark2Cure vary greatly in the difficulty and there is a large learning curve to become proficient at the task. Assuming that both factors affect the amount of time it takes for a user to completely mark a document and that users with more experience will completed the task at a more stable rate, we identified contributors to Mark2Cure who have annotated over 50 abstracts. We calculated their precision, recall, and f-score for their annotations with respect to the reference set. We also looked for successive submissions on a per user basis to estimate the time spent per submission and calculated the mean, median, minimum, and maximum amount of time (in minutes) spent per abstract. Abstracts which took longer than 45 min were presumed to have included some sort of break/distraction and were exclude from the time calculations. After obtaining user performance and time metrics, we binned the users by their f-scores and averaged the time metrics for each range/bin of f-scores.

### 2.7 Assumptions and limitations

Estimates are based on the simple assumption that the number of students enrolled in a school that has CSSLRs is the same as the number of students at the school who are impacted by CSSLRs are expected to provide ballpark figures for a snapshot in time. While student enrollment numbers in schools would ideally reflect the number of students served at those schools (and therefore subject to CSSLRs), the reality is more complicated as students may drop out, transfer, double-enroll, enroll part-time, graduate early, graduate late, etc.

## 3. Results

### 3.1 Number of students impacted and the hours of quality opportunities needed due to CSSLRs set at different levels of government

In the US, CSSLRs have been set at the state level (Maryland) and city level (Chicago) as well. Maryland high school students are required to complete 75 hrs of community service; while, high school students in Chicago are required to complete 40 hrs. In Canada, CSSLRs may be set at the province level such as in Ontario. As seen in Table 2a, requirements set at different levels of government can easily affect hundreds of thousands of students and create the need for millions of hours worth of opportunities a year.

**Table 2a.**
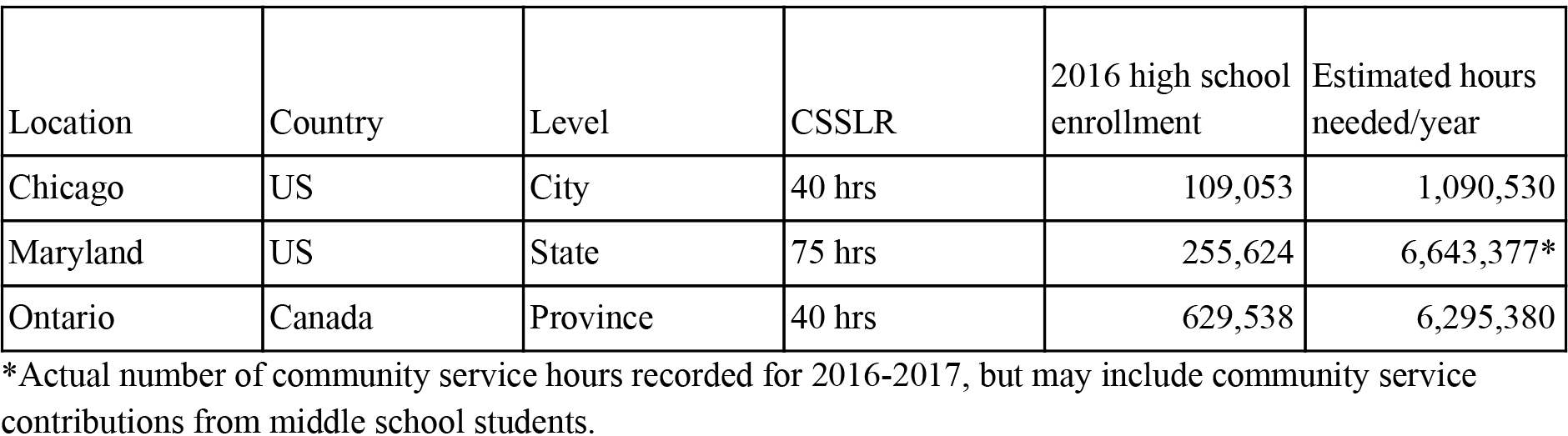
Examples of CSSLRs at different levels of government

In San Diego county, only two high school districts have district-wide CSSLRs: Sweetwater Union and Fallbrook Union. Outside of these two districts, some districts provided no restrictions allowing schools to impose their own CSSLRs. Even if a school did not have strict CSSLRs in place, there were student organizations and/or programs that have CSSLRs such as Key Club, National Honor Society, JROTC, and other programs that could be used for estimating the minimum number of students in need of volunteer hours at each school. According to data from the California Department of Education (Ed-data.org, 2018), there were 158,733 high school-aged students enrolled in public schools across San Diego county in the 2016-2017 school year. Of those students, 146,104 were enrolled in one of the public schools investigated in this study, while the remaining students were enrolled in schools excluded from the study due to the ‘non-traditional’ nature of the school (e.g., strictly online, or caters primarily to another age group.) One third of students in public schools across San Diego county (48,773) had a CSSLR to fulfill, and contribute 600,261 hours of community service annually. Across four years, these students will contribute an estimated 2.4 million hours of community service.

Even without a CSSLR in place, about 43% of the students in schools that collected data or provided estimates on student volunteerism engaged in community service activities. Assuming that students in participating in JROTC or Key club also engage in some form of community service, the number of San Diego County students in need of quality community service/volunteer opportunities rises to about 65,093. As seen in Table 2a and 1b, CSSLRs create a need for quality volunteer and community service opportunities. Citizen science participation may be able to meet those needs if the citizen science activities qualify as community service activities. To determine how citizen science organizations and efforts could fulfill CSSLRs, we investigated how CSSLRs and community service recommendations in San Diego County were tracked and/or validated.

**Table 2b.**
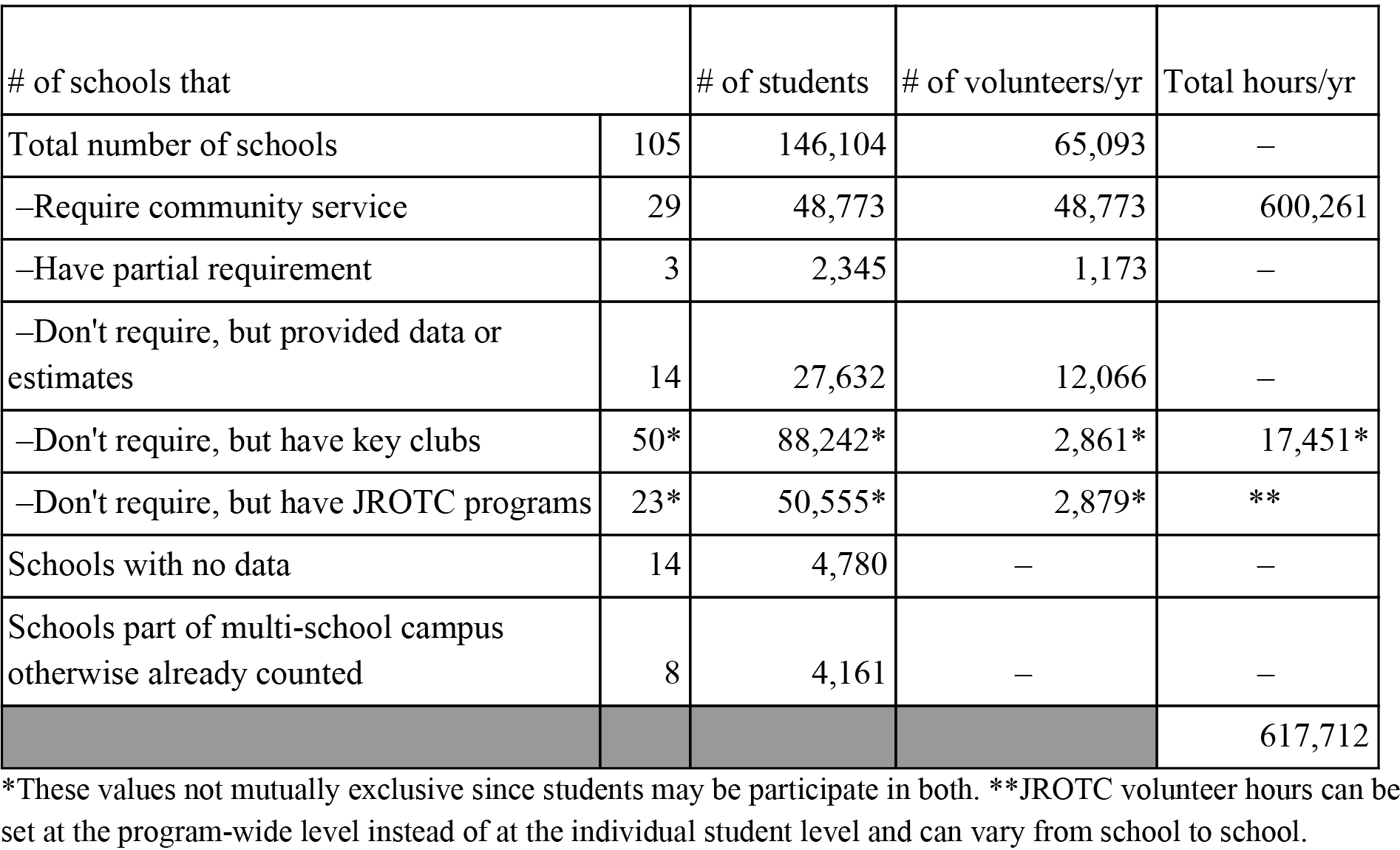
Summary of CSSLR impact on students in San Diego County in 2016-2017 academic year

| # of schools that |  | # of students | # of volunteers/yr | Total hours/yr |
| --- | --- | --- | --- | --- |
| Total number of schools | 105 | 146,104 | 65,093 | – |
| –Require community service | 29 | 48,773 | 48,773 | 600,261 |
| –Have partial requirement | 3 | 2,345 | 1,173 | – |
| –Don't require, but provided data or estimates | 14 | 27,632 | 12,066 | – |
| –Don't require, but have key clubs | 50* | 88,242* | 2,861* | 17,451* |
| –Don't require, but have JROTC programs | 23* | 50,555* | 2,879* | ** |
| Schools with no data | 14 | 4,780 | – | – |
| Schools part of multi-school campus otherwise already counted | 8 | 4,161 | – | – |
|  |  |  |  | 617,712 |

\*These values not mutually exclusive since students may be participate in both. \*\*JROTC volunteer hours can be set at the program-wide level instead of at the individual student level and can vary from school to school.

### 3.2 CSSLR and community service validation methods

Although only 29 out of the 105 schools had CSSLRs, only 24 of those 29 schools provided some sort of tracking form or some sort of guidance on the acceptable proof for meeting the CSSLRs. For 5 schools, no form or guidance could be found. For 3 of the 5 schools, the CSSLR was integrated (service learning) into the curricula. Although the majority of schools did NOT have a CSSLR, some schools still provided tracking forms on their websites, and tracked student volunteering activities in order to verify a student’s eligibility for various scholastic programs (e.g., Presidential Award, National Honor Society, CJSF, etc.) Of the schools that did provide some sort of means for tracking CSSLR fulfillment, most schools opted for some sort tracking form that could be signed off by a supervisor (or parent if necessary). Self-reporting trackers (no signature needed, but contact info should be included) were available from some schools without CSSLRs. CSSLRs were almost exclusively determined by time spent engaged in community service activities (minimum number of hours); therefore, citizen science efforts must be amenable to providing time metrics in order to engage participants fulfilling CSSLRs. To understand how citizen science efforts could align with the rationale behind CSSLRs, we analyzed the stated justifications and rationales that San Diego County high schools gave for requiring or encouraging community service activities and grouped them based on the Furco’s service program educational goals (Furco, 1994).

**Figure 1.**
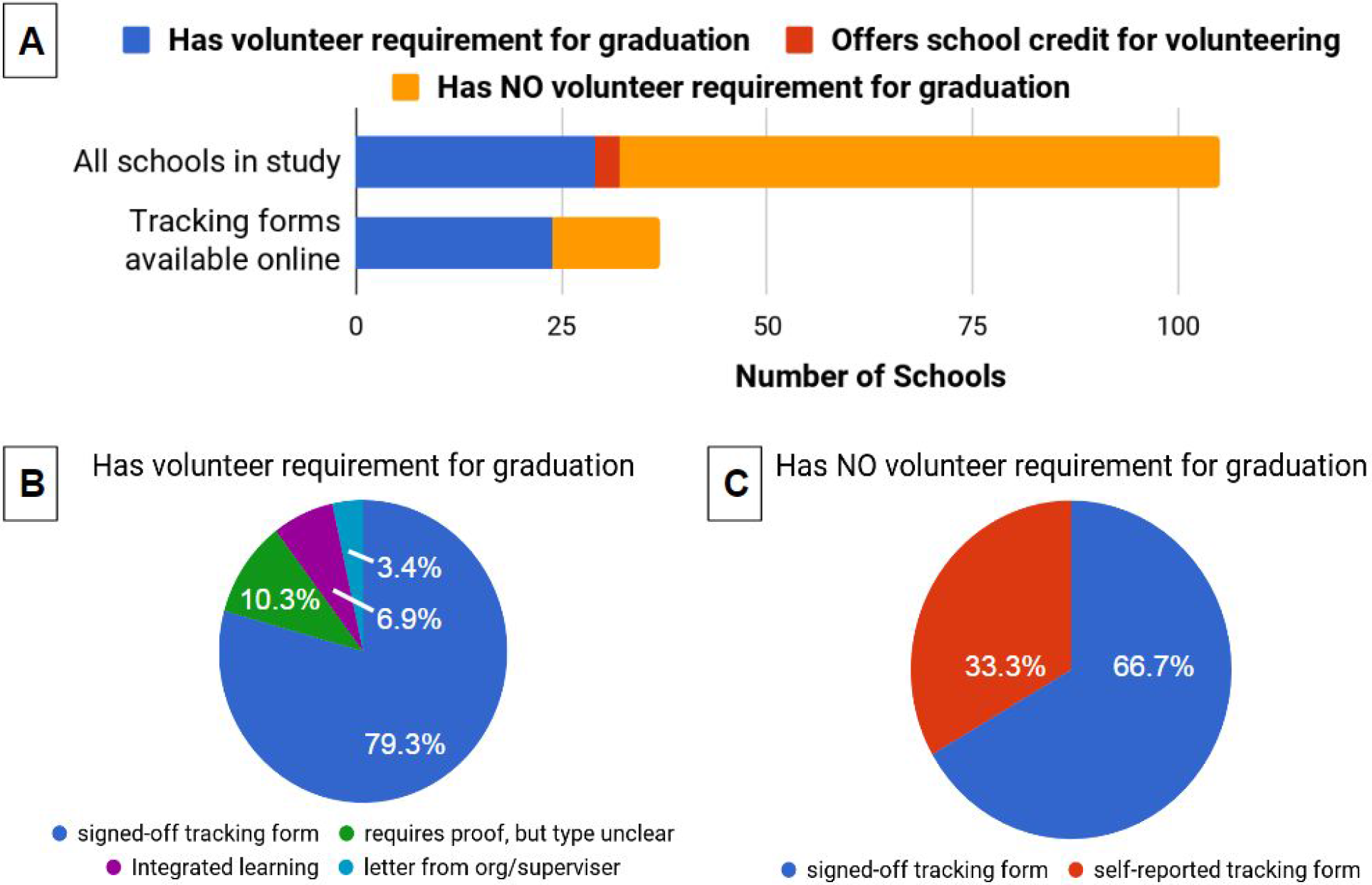
Volunteer requirements and acceptable proof. Number of schools with CSSLR, optional volunteering credits, or no requirements/credits as compared to schools with volunteering tracking forms online (A). Types of acceptable proof in schools with requirements (B) and those without requirements (C) based on the tracking forms available.

### 3.3 Stated justifications and rationales for CSSLRs or encouraging CSSL activities

Many different rationales were cited in support of CSSLR’s (Supp Figure 1B, Figure 2). Among San Diego County high schools with a formal requirement, the most widely-cited rationales were those which would encourage student’s social development. In contrast, schools which did NOT have CSSLRs, but recommended participation in community service activities tended to focus on academic achievement justifications. While some schools do have guidelines in place as to what counts as a community service activity, guidelines are still subject to interpretation. To make it easier to understand how a citizen science project could serve to fulfill a CSSLR, citizen science projects should tailor their messaging to reinforce the rationale for the community service recommendation / requirement. Based on these findings, the messaging in San Diego County should focus on college admissions (academic achievement) and the public benefit (social development) if appealing to schools with no CSSLRs in place. For San Diego County high schools with CSSLR obligations, messaging should focus on public benefit and relationship building/community engagement (social and political development).

**Figure 2.**
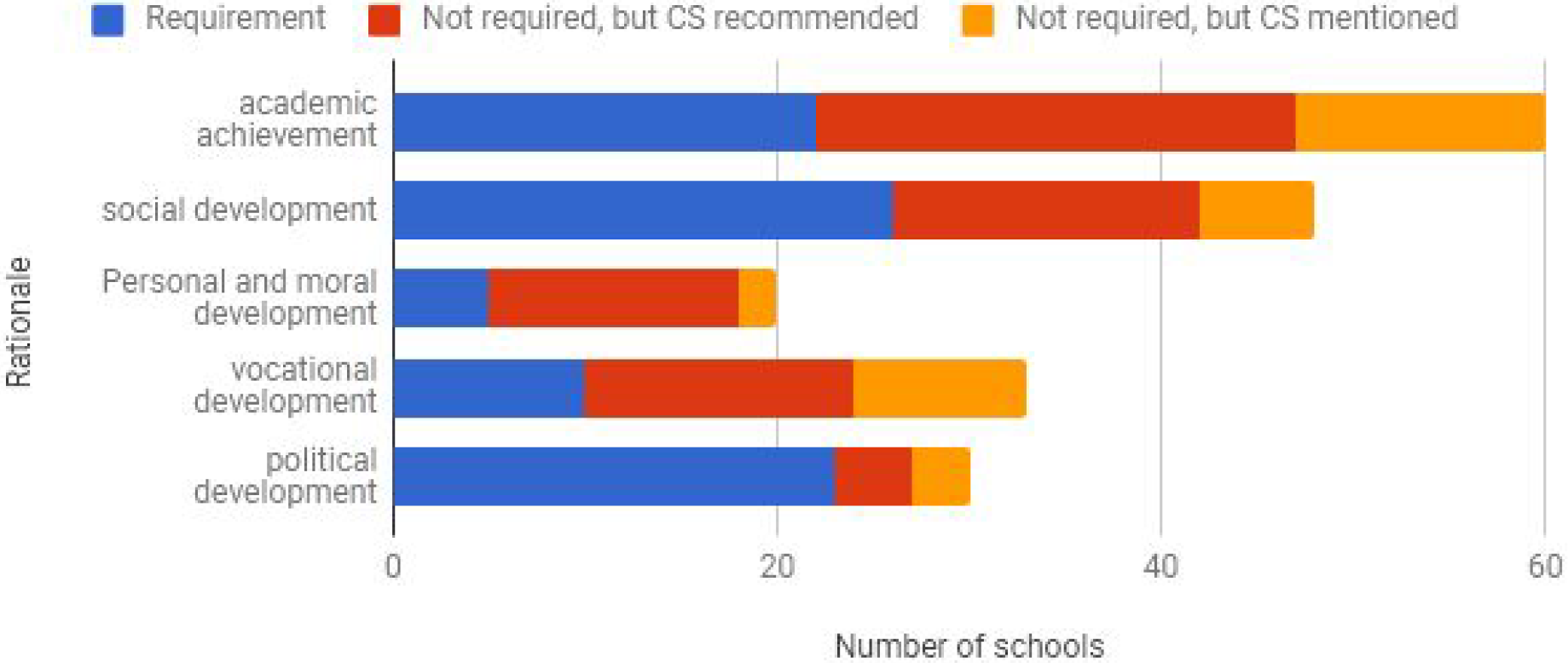
Educational rationale for CSSL activities provided by schools in San Diego County. The frequency of a particular educational category of rationale cited in support of schools that had CSSLRs (blue), did not have CSSLRs but recommended community service (red), and did not have CSSLRs but mentioned community service (orange).

To understand the criteria needed for a citizen science project to qualify as a community service activity, we analyzed the definitions, examples, and areas of impact of community service provided by schools with CSSLRs. We identified 101 health and medical citizen science projects with the assumption that these projects were more likely to impact human health and therefore qualify as a community service activity, and reviewed these projects in terms of their suitability for CSSLR fulfillment based on the qualification and validation criteria used by San Diego high schools with CSSLRs.

### 3.4 Definitions of community service–basic qualifications for citizen science projects to meet

Many schools encouraged students to engage in community service and even provided verification forms; however, actual guidelines on what counted as community service were fewer in number and tended to come from schools with CSSLRs. Only six unique examples/guidelines on community service activities were found. The non-community service (Not CS) and community service (CS) activities from the six examples were aggregated as seen in Supplemental Table 1.

In general, an activity counted as community service if it meets all of the following criteria:

1. Activity is done under adult supervision of someone outside of the family / Activity can be verified by someone outside of the family
2. Activity is done in conjunction with non-profit organization that benefits community
3. Activity is done without payment
4. Activity is done outside regular school hours or typical extracurricular activities (such as school sports fundraising)
5. Activity actually helps others, is not just a meeting or talking about helping.

The above criteria for CSSL activities derived from San Diego County High Schools with CSSLRs were consistent with those from Chicago Public Schools (Chicago Public Schools, n.d.) and the state of Maryland (Maryland State Department of Education, 2017).

### 3.5 Suitability of health and medical citizen science projects in SciStarter for fulfilling CSSLRs

**Table 3.**
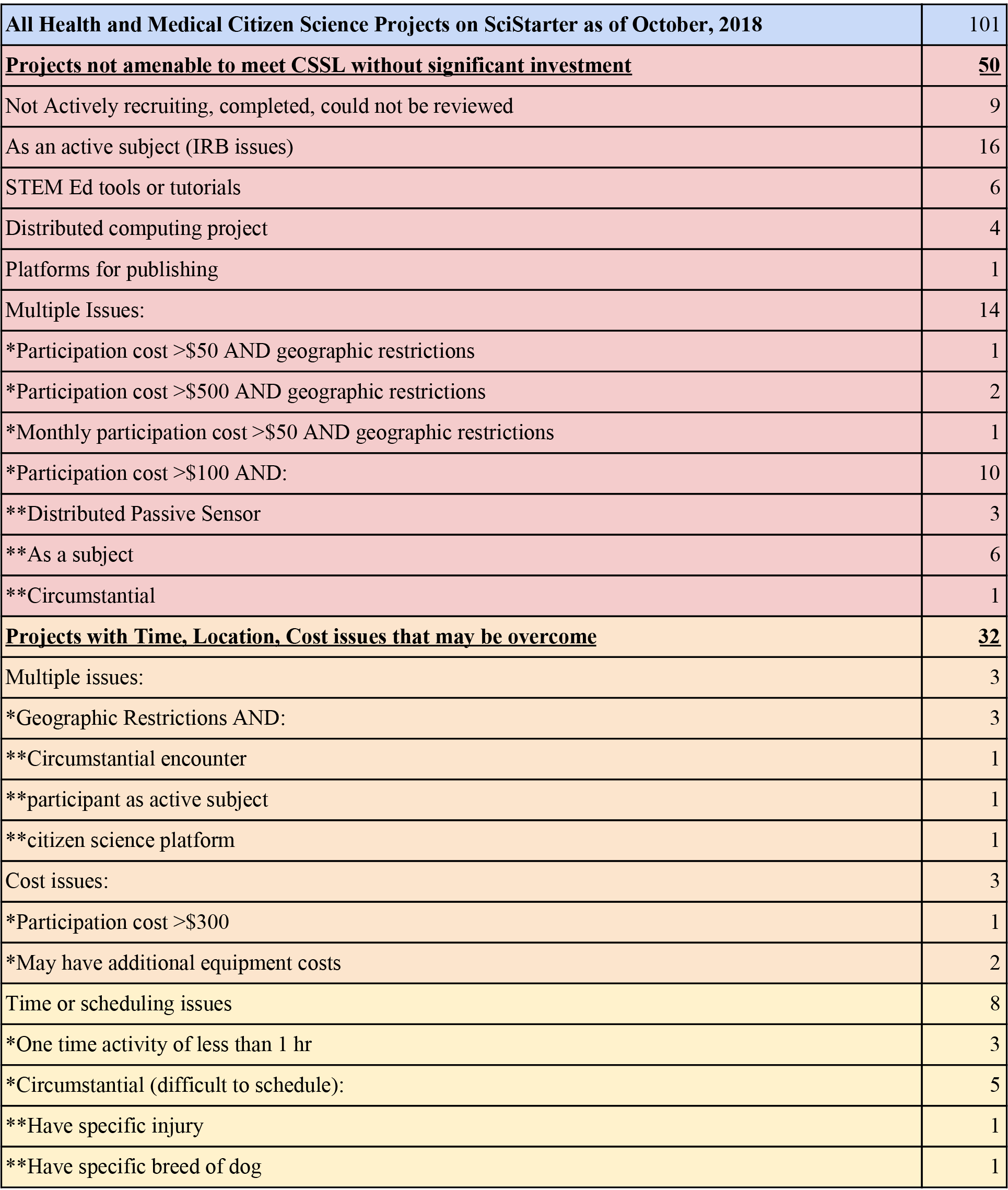
Compatibility issues with health and medical citizen science projects from SciStarter. Red indicates issues which render a project incompatible with CSSLRs including ethical and legal issues, project status, limitations on active participation, and for profit status of project organizer. Yellow and orange indicate time, cost, location issues which may render a project more difficult for students of lower socioeconomic status. Green indicates projects which do not have these issues.

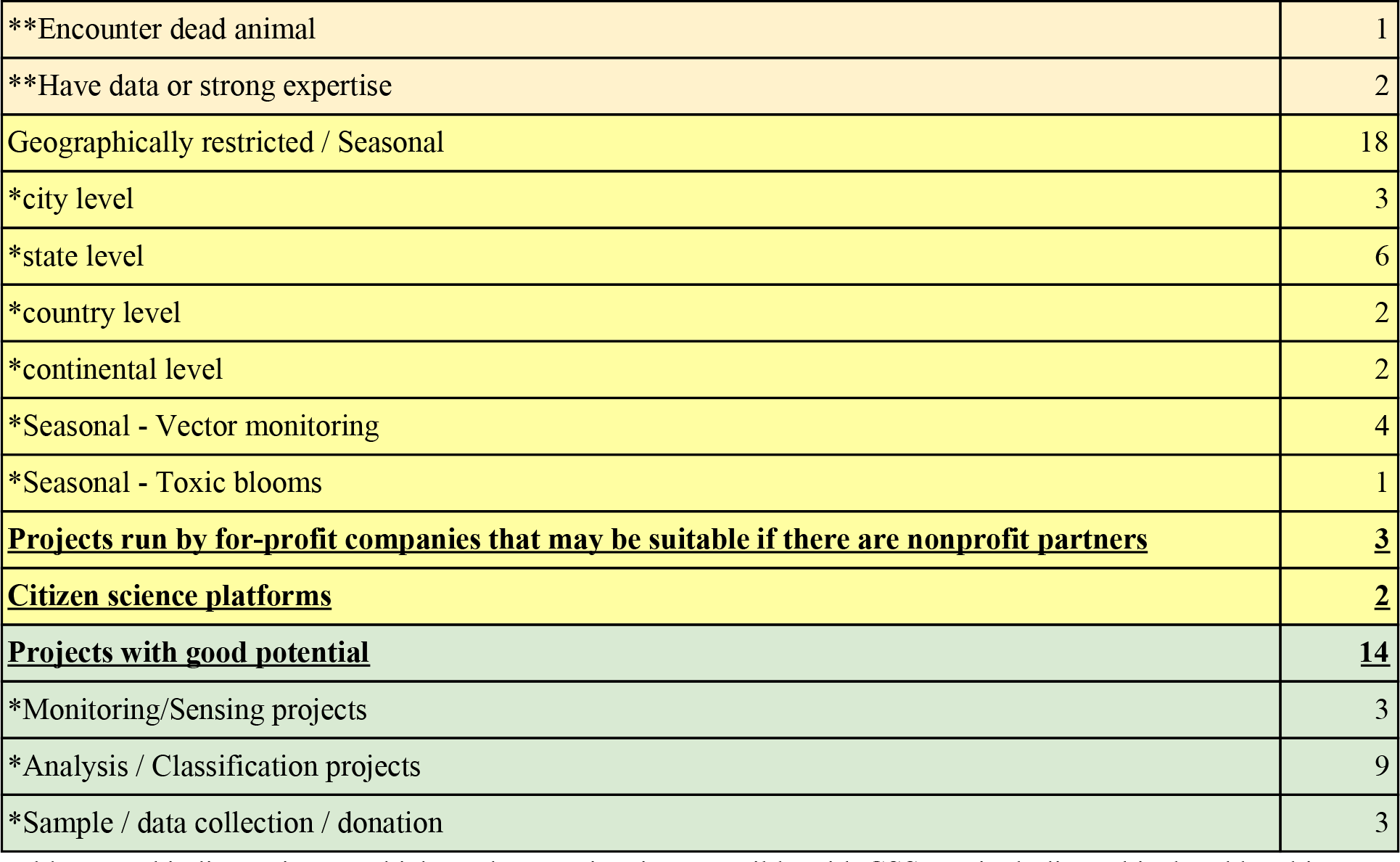

Of the 101 citizen science health / medical citizen science projects identified from SciStarter, 50 projects were deemed not amenable to meet CSSL without significant investment. Of those 50, nine projects were either not actively recruiting, completed, or could not be reviewed because the site was down. Six projects appeared to be tools for STEM education or tutorials for citizen science projects--but not the actual project, four were distributed computing projects (in which volunteers leave their computers running a program and are otherwise not engaged) and one was a self-described open publishing platform. In 16 projects the volunteer collects and donates information on himself or herself effectively becoming both a citizen scientist and a research subject. As we are focused on students fulfilling CSSLRs most of these students will not be old enough to consent to participating as research subjects. Fourteen projects had multiple issues which would rendered them difficult to use for CSSLs without significant investment of time and money. For example, six projects would cost over $80 to participate as an active subject (legal/ethical issues), and three projects involved the purchasing and installation of a monitor (which can automatically send data and can disengage the contributor).

Thirty two projects had time/cost/location issues which unduly impact students of lower socioeconomic status. Three projects had multiple issues which could be overcome by chance/circumstance. Three projects had cost issues, 8 projects had time issues or were heavily circumstantial (which could be difficult to schedule or could require more time or money to overcome), and 18 projects had location issues and were geographically and/or seasonally dependent. These projects would be suitable for students within specific geographic and/or seasonal boundaries who have means of transportation as required by some of the projects. Two “projects” were actually citizen science platforms from which an appropriate project might be found with additional work. Three projects were tied to for profit companies which could disqualify them as a CSSL activity if they do not have non-profit partnerships (at least one had a nonprofit partnership).

Fourteen online or location indifferent citizen science projects were devoid of the aforementioned issues and would likely be good activities for fulfilling CSSLRs, namely: Project Soothe, Field-Photo-Library, MalariaSpot, Colony B, EteRNA, ANT-vasion, AgeGuess, Flu Near You, Globe at Night, Hush City, EyeWire, Cochrane Crowd, Stall Catchers, and Mark2Cure. To illustrate how online / location indifferent citizen science projects may need to be adjusted for students fulfilling CSSLRs, we align the contributions metrics of our own project, Mark2Cure with the verification requirements of CSSLRs.

### 3.6 Aligning contribution metrics in Mark2Cure with CSSLRs

Mark2Cure is a citizen science project aimed at organizing biomedical knowledge to facilitate disease and treatment research to ultimately improve patient outcomes. Although helping patient populations is an acknowledged community service activity and naturally aligns with Mark2Cure, Mark2Cure is an online platform and does not readily suit the needs of students in need of community service opportunities. Schools generally measure community service contributions in terms of hours, while virtual citizen science projects like Mark2Cure tend to measure contributions in terms of classifications, annotations, submissions, etc. In order to empower students to fulfill their obligations while contributing to a citizen science project like Mark2Cure, we need to align our contribution metrics with those acceptable by schools and provide an easy way of verifying their efforts.

Volunteer and citizen science contributions through Mark2Cure are time stamped, allowing successive contributions from the same user to be used to estimate the time that user spent on a task. The use of this timestamp to estimate the time a contributor spends on an task is problematic for two reasons:

1. It can be very inaccurate. Estimating time spent on a task based on the difference between timestamps of a contributors successive submissions will include time they simply left the task open.
2. It does not reflect the true value of the contributor’s submission. A contributor’s submission is most valuable when they try to do the task correctly.

Another solution would provide a simple conversion factor to convert number of contributions into time; however, this conversion factor could incentivize students to work quickly without regard for quality. To correct for this, we analyzed our contributions to create a scaled conversion factor.

On average users performing at a level of 0.75-0.8 spend a median of 4.4 minutes per abstract (Figure 3). User performing at even higher levels (>0.8) spend even less time per abstract, presumably because increased expertise decreases the time needed to perform a task well. Based on this analysis, Mark2Cure could apply a scaled conversion factor that would reward students to perform at higher levels. For example, each abstract completed at a level of 0.75 or above would count at 5 minutes of volunteer time. Abstracts completed at a level of 0.7 to 0.75 would count as 3 minutes of volunteer time; while, abstracts completed at a level of less than 0.7 would count as 2 minutes of volunteer time. In this manner, students are incentivized to contribute higher quality annotations over lower ones.

**Figure 3.**
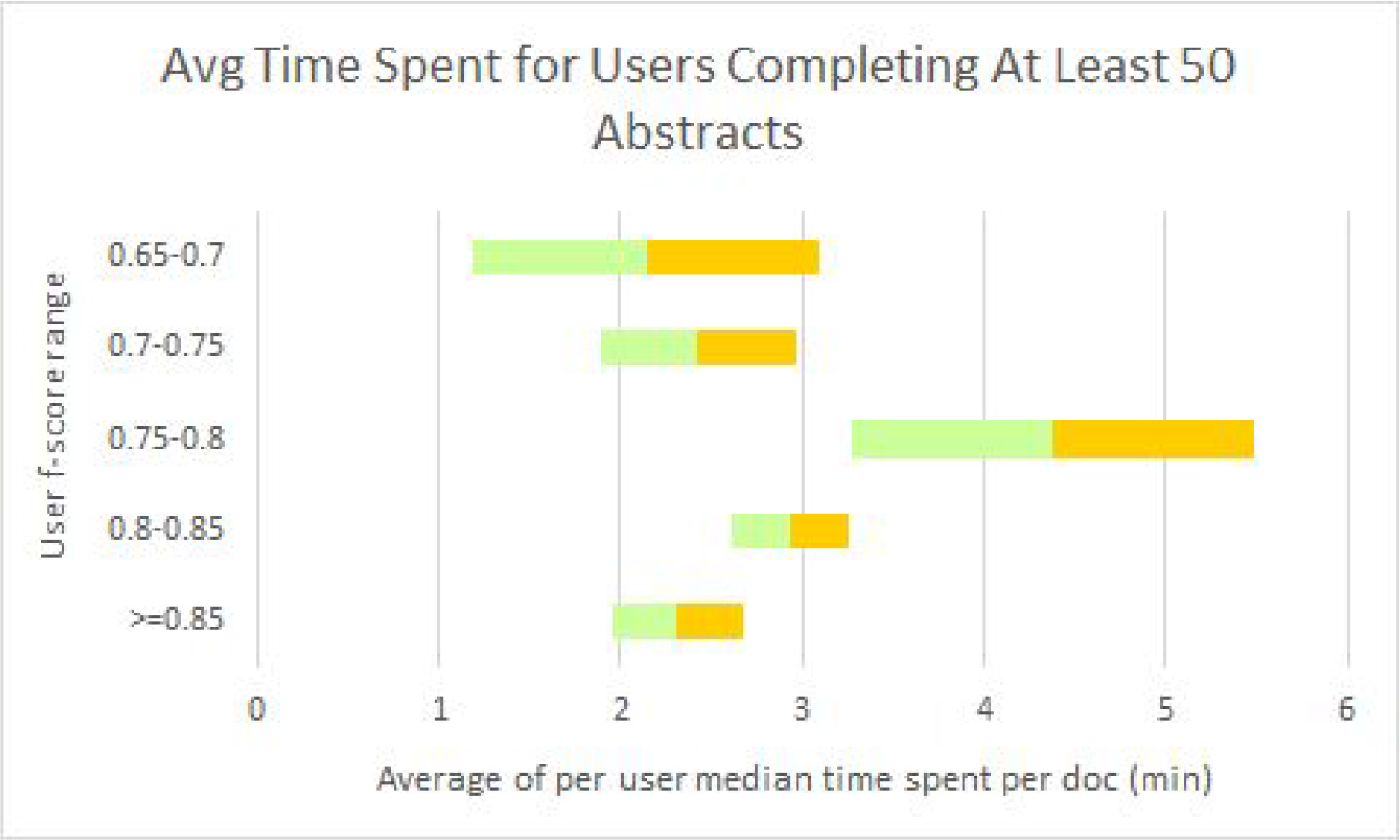
Time vs NER quality in Mark2Cure. The average time spent per abstract is represented by the line between the green and orange bars which represent the lower (green) and upper (orange) standard error of the mean.

To ensure that these conversion factors were reasonable, we selected two high school students who contributed annotations on over 100 abstracts and calculated their individual performance and time metrics. Both students had f-scores of above 0.8 relative to the reference set. The median amount of time spent per abstract by each of the students was 1.6 and 3.6 minutes–averaging 2.6 minutes, which is within the lower standard error of the mean for the corresponding user f-score range. Users with average f-scores below 0.65 relative to the reference set generally did not contribute at least 50 abstracts and were not included.

As an online platform, the conversion of Mark2Cure metrics and population of a CSSLR log/form could potentially be automated. To determine the fields that would be important for such a process, we documented the fields available in the 25 different CSSLR logs/forms/trackers found from school websites in San Diego County (Table 4). Schools in districts with CSSLRs were assumed to use the district-wide forms which were counted only once for each district. The most common fields on the verification trackers/forms/logs provided to these students were the Student’s name, the name of the sponsoring organization, the location of the community service activity, a description of the activity, the date the student participated in the activity, the number of hours spent on the activity, the supervisor’s name, contact info, and signature. Some forms requested more granular details about the student, organization, activity, or supervisor.

**Table 4.** Verification form field frequencies

| Data Category | Tracking form data fields | Totals |
| --- | --- | --- |
| School | School Preauthorization required? | 5 |
|  | Parent Preauthorization required? | 4 |
|  | Teacher/Advisor/school rep name | 4 |
|  | club/org | 2 |
| Student | Student name | 25 |
|  | Student ID | 11 |
|  | Student expected year of graduation | 5 |
|  | student grade | 8 |
|  | school year | 4 |
| Organization | Service site / sponsoring org name | 19 |
|  | Business card or letter on letterhead | 2 |
|  | sponsoring org website | 3 |
|  | Location / Place of service | 12 |
| Activity | Day / Date | 24 |
|  | Event | 2 |
|  | Activity description | 23 |
|  | time start | 9 |
|  | time end | 9 |
|  | Number of hours | 25 |
| Supervisor | supervisor name | 19 |
|  | supervisor contact info | 19 |
|  | supervisor position | 7 |
|  | Supervisor signature | 21 |
|  | Supervisor initials | 2 |
|  | supervisor comments | 2 |
| Reflection | Reflection on activity | 2 |

### 3.7 Recommendations for aligning citizen science projects with CSSLRs

Based on the results of our investigation, we provide the following recommendations to citizen science projects for aligning with CSSLRs:

1. As discussed in the introduction, community service is volunteerism that benefits the community, service learning integrates community service activities with course curricula to improve the value of the service learning to the student. For this reason, citizen science project goals should be framed in terms of its value to the community (i.e.-the ‘big picture’ public good) as well as its value to the student (e.g.-engaging the student with the community, or the value of these experiences for college applications). If the justifications for CSSLRs in a specific region are known, citizen science projects hoping to recruit from that region should align the benefits of their project with the rationale given for having CSSLRs in that region. For example, mycological biodiversity efforts in San Diego County may want to emphasize the value of a student’s contributions to this scientific endeavor, the importance of fungi to ecological health, and the impact of ecological health has to a community’s economic and physical wellbeing.
2. In San Diego County, CSSL activities had to be performed in conjunction with a non-profit organization that benefited the community in order to count (section 3.4), thus citizen science projects should associate with the non-profit entities that sponsor them. If a citizen science project is born under a for-profit and nonprofit partnership–the nonprofit association should be emphasized.
3. All 25 of the San Diego County high school CSSLR tracking forms included a field for recording the time spent on an activity (Table 4). CSSLRs imposed at different levels of government were all imposed as a temporal requirement (Table 2a); hence, citizen science projects need to document and provide time/temporal feedback. CSSLRs are almost exclusive time-dependent requirements, so projects that forsake time measurements for quality metrics must find a way to accommodate.
4. 96% of the San Diego County high school CSSLR tracking forms included a field for organizational supervisor signature (Table 4). Citizen science projects should have a team member or coordinator available for answering questions from students, tracking/verifying student participation, and providing necessary validation (such as signing forms).
5. Citizen science projects need to be findable by students seeking to fulfill CSSLRs. Citizen science projects should contact local schools to get listed on school websites (if available), and be added to volunteer listing sites/matching services like VolunteerMatch.org and AllForGood.org.

## 4. Discussion

Citizen science is already an acknowledged tool for ISE (Shirk et al. 2012) and practitioners have been actively investigating its integration in FSE. The gap between ISE and FSE can be difficult for citizen science project leads to overcome without sufficient resources. For citizen science to successfully make a transition from ISE to FSE, citizen science organizations will need to establish partnerships with STEM educators interested in the co-creation of educator resources. Research from the science education field suggests a middle ground (Non-Formal Science Education, NFSE) which could bridge the gap. This middle ground, NFSE, is described as typically having some structure, being prearranged, usually voluntary (thus primarily being intrinsically motivated, but not always), occurring at an institution outside of schools (such as museums, zoos, aquariums, etc), and where learning is not usually evaluated strictly (Eshach, 2007). The characterization of NFSE is primarily used to explore science learning in the context of museum/zoo field trips or mobile museums/zoos, but could just as easily describe CSSLRs applied to a scientific endeavor. Hence, the fulfillment CSSLRs with citizen science participation could be considered a form of NFSE and serve as an important bridge for projects interested in FSE integration.

Volunteerism has been extensively studied in the context of education and service learning. The merits and disadvantages of having CSSLRs (mandating volunteerism) in schools is subject to ongoing research and debate (Helms, 2013)(Kim and Morgül, 2017)(Bode, 2017). In general many of the benefits of well implemented CSSLRs have been confirmed; however, good implementation takes time, resources, and effort. While schools iteratively improve their CSSLR implementation over time, it is the students and the educators who bear the burden of the gaps in support in the interim. For these students and educators, citizen science may provide opportunities for fulfilling CSSLRs in spite of the gaps in support similar to how CSSLRs may serve as an important entry point for citizen science projects lacking resources to bridge the gap between ISE and FSE. Citizen science projects, especially online citizen science projects, may be particularly useful to students (especially students of lower socioeconomic status) with CSSLRs to fulfill in the face of time, cost, and location barriers. While citizen science projects may serve as good opportunities for fulfilling CSSLRs for some students, differences in technological access may limit the number of virtual citizen science projects that could be leveraged to fulfill a CSSLR. In spite of dramatic improvements in providing internet access in public schools (Vigdor, Ladd and Martinez, 2014) and libraries (Horrigan, 2016), many students still lack traditional broadband access at home (Lei and Zhou, 2012). Fortunately, many virtual citizen science projects are also mobile-friendly and smartphone/ cellular technologies have begun to fill the internet access needs for people lacking home access to traditional broadband in urban and suburban communities (Smith, 2015)(Vangeepuram et al., 2018). In rural communities, the digital divide persists due to infrastructure issues; however, there many citizen science projects involving the environment and nature that require only occasional online access and are suitable for participation in geographically remote areas.

## 5. Conclusions

Given the natural intersections between the fields of citizen science, STEM education, and volunteerism, CSSLRs present an interesting area of investigation. We inspected the prevalence of CSSLRs in San Diego County and at different levels of government and estimated the potential market need created by CSSLRs. We analyzed the rationale and verification methods used for those CSSLRs and provided guidance on how citizen science efforts (especially online citizen science efforts) can be tweaked to meet them using own platform, Mark2Cure, as an example. In the spirit of citizen science and to extend our knowledge beyond the data we were able to collect in this research, we have added our San Diego County CSSLR data to citsci.org (http://citsci.org/cwis438/browse/project/Project_Info.php?ProjectID=2108) and created the fields needed in order for anyone to collect and contribute data on CSSLRs in their local schools. We hope this data will be useful to citizen science organizations/platforms like Scistarter.com which are becoming an increasingly valuable resource for STEM educators. CSSL tracking form data, CSSLR mentions in San Diego County High Schools, and the SciStarter Biomed/Health project analysis can be found at: https://github.com/gtsueng/CSSLR-data.

## 6. Acknowledgments

We thank VolunteerMatch for connecting us to Arun (one of the authors on this paper) an international business student who applied his market research background towards surveying the regional service requirements. We thank our many Mark2Curators for their contributions, criticisms, feedback, and suggestions. The list of Mark2Curators who contributed to the data set used in this paper and permitted us to publish their names can be found at: https://mark2cure.org/blog/csslr-paper-contributors/. We thank the students Kathryn Jin, Nina Lu, and Celine Son sharing their experiences and contributing to Mark2Cure. We also thank the many counselors, principals, and administrators in high schools across San Diego County who took the time to answer our inquiries. We thank SciStarter for making it easy to quickly identify biomedical and health citizen science projects.

Mark2Cure is supported by the US National Institute of Health (U54GM114833 and GM089820 to A.I.S.) and by the Scripps Translational Science Institute, an NIH-NCATS Clinical and Translational Science Award (CTSA; 4 UL1 R001114).

