## Supplemental Figure and Table for "Aligning Needs: Integrating Citizen Science Efforts into Schools Through Service Requirements"

### Supplemental Figures and Tables

Supplementary Figure 1- Sample binning of Community Service mentions

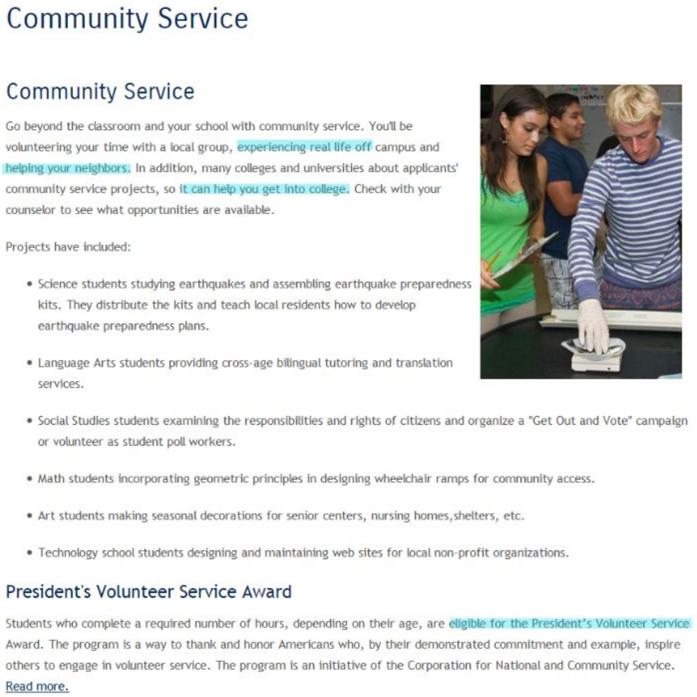

#### Community Service

##### Community Service

Go beyond the classroom and your school with community service. You'll be volunteering your time with a local group, experiencing real life off campus and helping your neighbors. In addition, many colleges and universities about applicants' community service projects, so it can help you get into college. Check with your counselor to see what opportunities are available.

Projects have included:

- Science students studying earthquakes and assembling earthquake preparedness kits. They distribute the kits and teach local residents how to develop earthquake preparedness plans.
- Language Arts students providing cross-age bilingual tutoring and translation services.
- Social Studies students examining the responsibilities and rights of citizens and organize a "Get Out and Vote" campaign or volunteer as student poll workers.
- Math students incorporating geometric principles in designing wheelchair ramps for community access.
- Art students making seasonal decorations for senior centers, nursing homes,shelters, etc.
- Technology school students designing and maintaining web sites for local non-profit organizations.

##### President's Volunteer Service Award

Students who complete a required number of hours, depending on their age, are eligible for the President's Volunteer Service Award. The program is a way to thank and honor Americans who, by their demonstrated commitment and example, inspire others to engage in volunteer service. The program is an initiative of the Corporation for National and Community Service.
Read more.

| | |
|---|---|
| 1 | Because college admissions |
| 0 | Foster civic responsibility and community/engagement, relationships |
| 0 | Learn/buildskills |
| 1 | improvecommunity/school/help neighbors |
| 0 | build esteem, good feels, character |
| 1 | real life experience |
| 1 | qualify for award, association, guaranteedadmission, IB qualification etc |
| 0 | for fun/interest/passion |
| 0 | class requirement |
| 0 | Intervention/ penalty |

Supplementary Figure 1B - Expressed justifications and rationale for community service

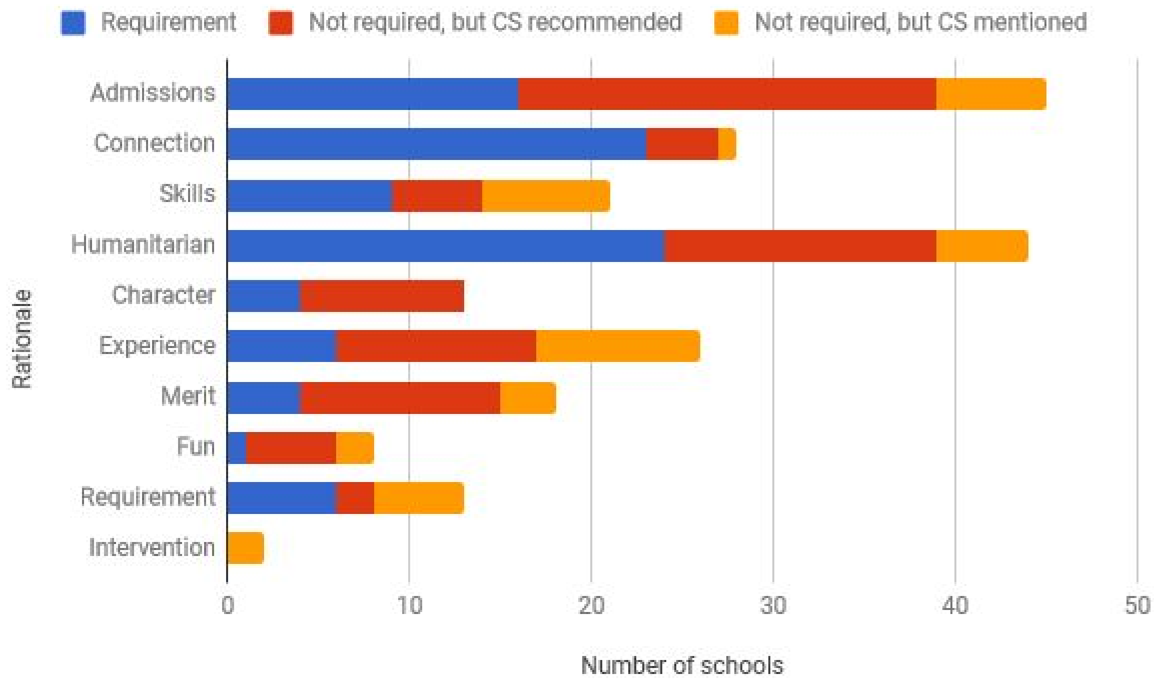


The frequency of a particular category of rationale cited in support of schools that had CSSLRs (blue), did not have CSSLRs but recommended community service (red), and did not have CSSLRs but mentioned community service (orange). Admissions were mentions of college admissions.  Connection indicated motivations of connecting the student to the broader community such as fostering civic responsibility and community/engagement, relationships. Skills included learning, applying, or building skills. Humanitarian reasons were those involving improve community, improving the school, helping neighbors and people in need. Character included building esteem, feeling good about oneself, and building character. Experience was just mentions of gaining real life experience. Merit included mentions of qualifying for an award, acceptance into an association, guaranteed admission, IB qualification etc. Fun included fun, hobby, interest, passion. Requirement is NOT the same as schoolwide or club/organization based CSSLRs, but instead refers to community service requirements set for a particular or specific class not necessarily available to all students (e.g., AVID). Intervention includes justifications of community service as a penalty for misbehavior.

Supplemental Table 1 - Aggregate List of Activities that Qualify or Don’t Qualify as Community Service Activities

| Sample Activity | Not CS | CS |
|---|---|---|
| Paid work | 4 | 0 |
| Sales of items for fundraisers for school or sports | 4 | 0 |
| Attendance in club meetings, church, youth group | 3 | 0 |
| Service done for family members or neighbors | 4 | 0 |
| Service related to a class, credit for a class, or the making of profit, defraying costs of trips, etc. | 2 | 0 |
| Service performed for a profit-making organization | 2 | 0 |
| Activities that would usually be considered normal extracurricular (or co-curricular) activities such as sports and sports related activities (managers), cheerleading, participating in school performance activities that are related to a class, ASB activities, etc. | 2 | 0 |
| Service for Hospital, soup kitchen, homeless shelters, or orphanage | 0 | 3 |
| Service for Non-profit, community-based, professional organization | 0 | 3 |
| Animal rescues | 0 | 2 |
| Assisted Living Facilities, convalescent homes | 0 | 2 |
| Community service through a religious institution | 0 | 2 |
| Community service club activities (not meetings) | 0 | 2 |
| Community service through Boy Scouts or Girl Scouts | 0 | 2 |
| Organized Community Clean up / beautification activities | 0 | 2 |
| Political campaign activities | 0 | 2 |
| Tutoring outside of school hours | 0 | 2 |
| Sports events of younger children, refereeing, etc. | 0 | 2 |
| Unpaid poll worker on Election Day (must be organized through school) | 0 | 2 |

Numbers do not add up to six since each example/guideline did not have every activity listed. For example, four of the six unique examples/guidelines found specifically mentioned that paid work would not count as a community service activity. The remaining two examples/guidelines made no mention of paid work.
